## supplementary_figures for "Large-scale network analysis captures biological features of bacterial plasmids"

**Acman et al.**

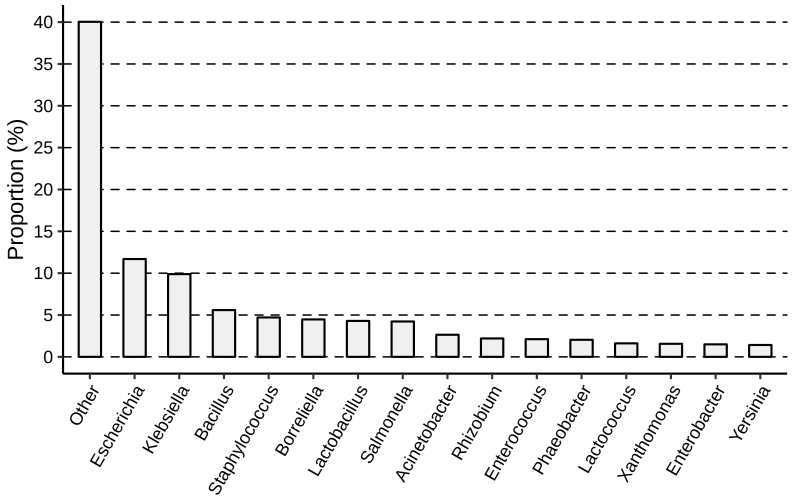


**Supplementary Figure 1. Phylogenetic diversity of plasmid hosts at the genus level.**

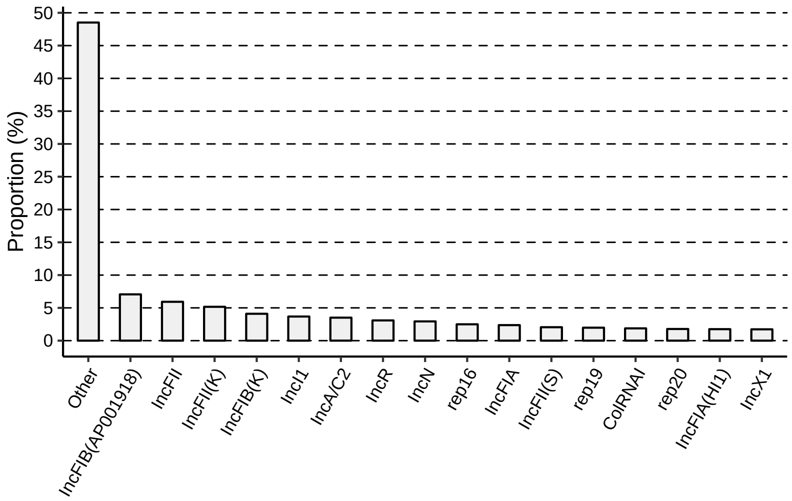


**Supplementary Figure 2. Distribution of the proportion of known plasmid replicon types.**

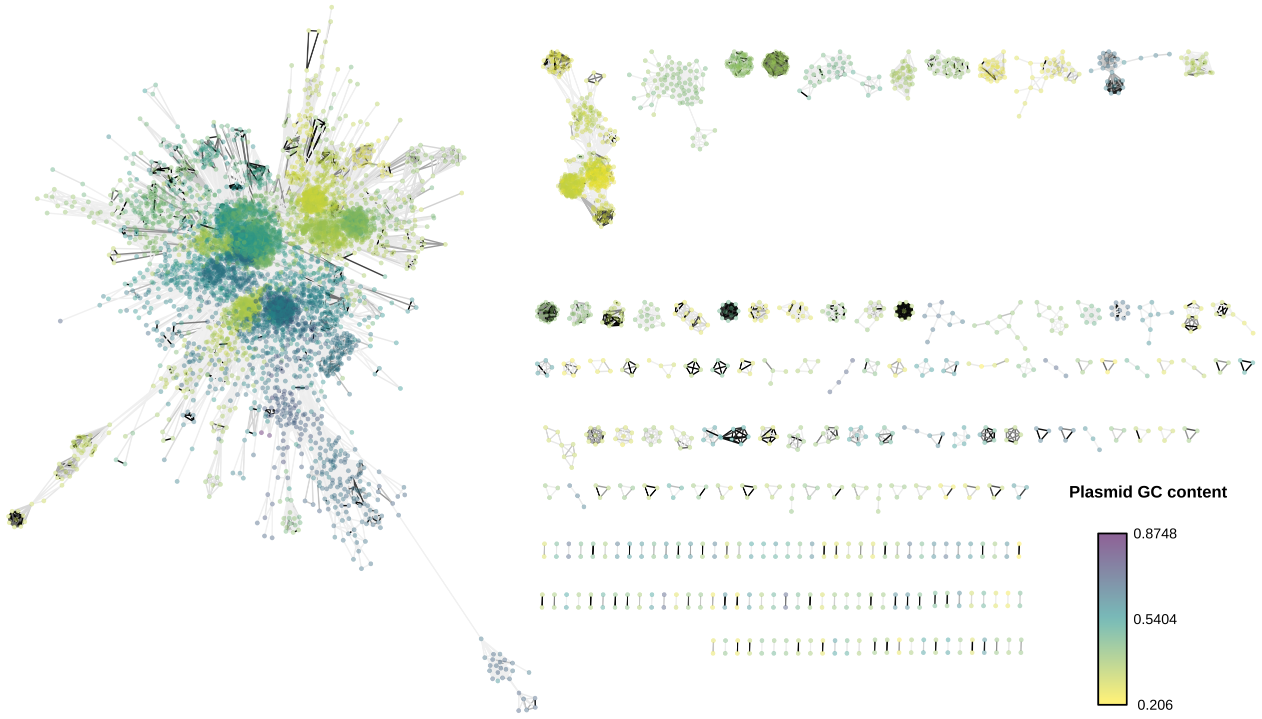


**Supplementary Figure 3. A network of plasmids (network density = 0.0438).** 10,696 plasmids (vertices) are connected by weighted edges where grey-scale colour gradient specifies the Jaccard Index (JI) similarity. JI is calculated as the proportion of shared *k*-mers between pairs of plasmids with a darker shade indicating higher similarity. Plasmid pairs which share less than 100 *k*-mers are considered to have a JI equal to zero. The colour gradient of the nodes indicates the GC content of plasmids. This representation depicts clustering of plasmids according to their GC content and hints at an underlying population structure.

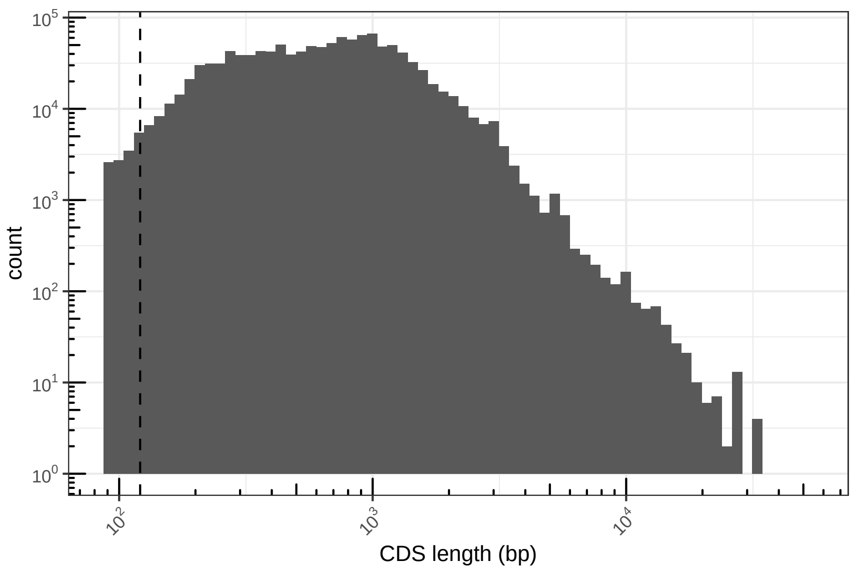


**Supplementary Figure 4. The distribution of the lengths of the plasmid-borne coding sequences (CDSs).** Both axes are in logarithmic scale. The vertical dashed line represents the cut-off value (<100 *k*-mers) applied while calculating JI similarity between plasmid pairs. Since the length of *k*-mers used was 21bp, the effective cut-off value applied is 121bp. Thus, few CDSs shared between plasmids with length less than 121bp may have been missed with the implementation of this cut-off.

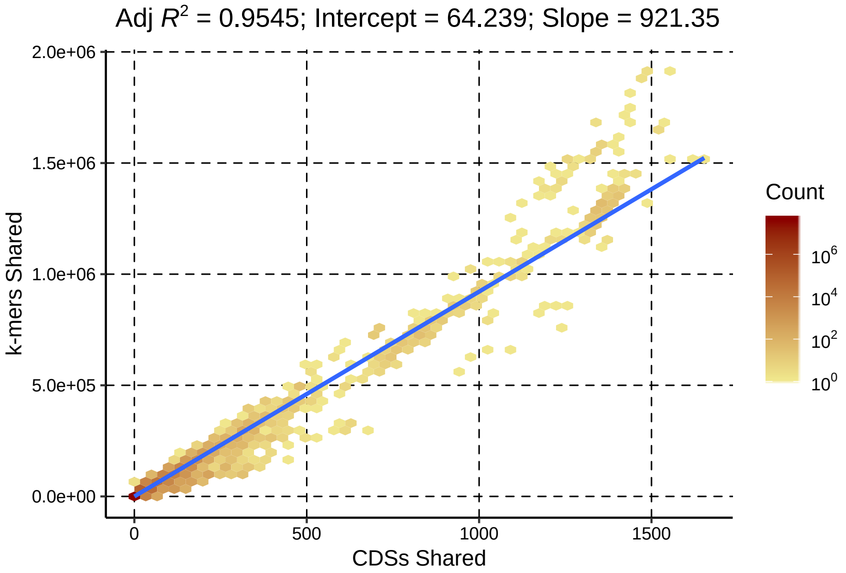


**Supplementary Figure 5. Linear correlation between number of shared CDSs and number of shared *k*-mers in plasmid pairs.** To facilitate the visual interpretation, the data points were grouped into hexagonal areas of equal size. The colour intensity of each hexagon reflects the density (count scale) of the data points in that particular area. An intercept (64.239) of the regression line (blue) suggests that a plasmid pair sharing around 64 *k*-mers on average do not have any CDSs in common.

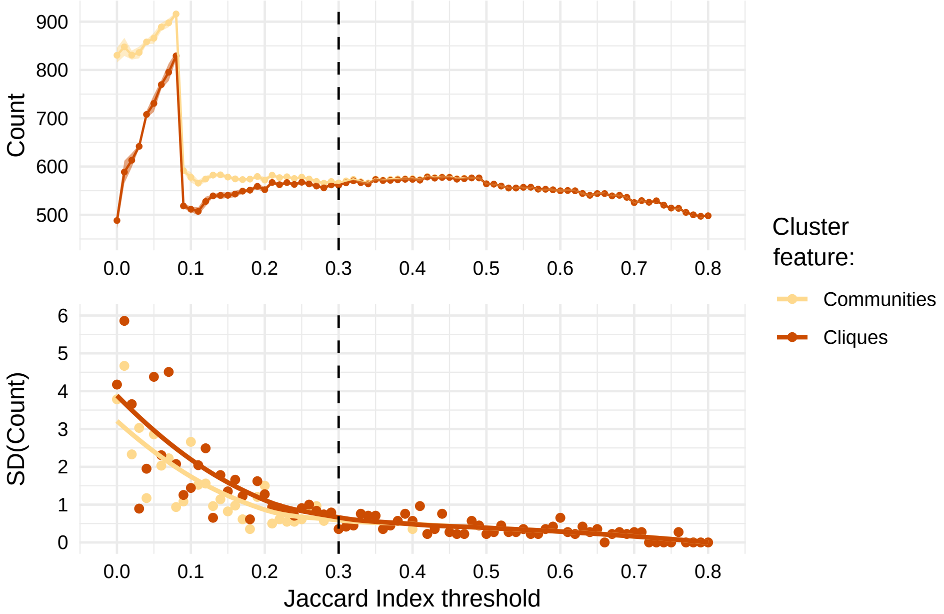


**Supplementary Figure 6. Optimization of OSLOM performance.** Additional criterion in determining the optimal JI threshold by assessing the consistency of OSLOM community detection performance. The upper plot shows the number of communities and cliques detected by OSLOM for each tested JI threshold. The lower plot depicts the drop in standard deviation (SD) of the number of cliques and communities detected as the network becomes sparser. The dashed vertical line represents the 0.3 JI threshold.

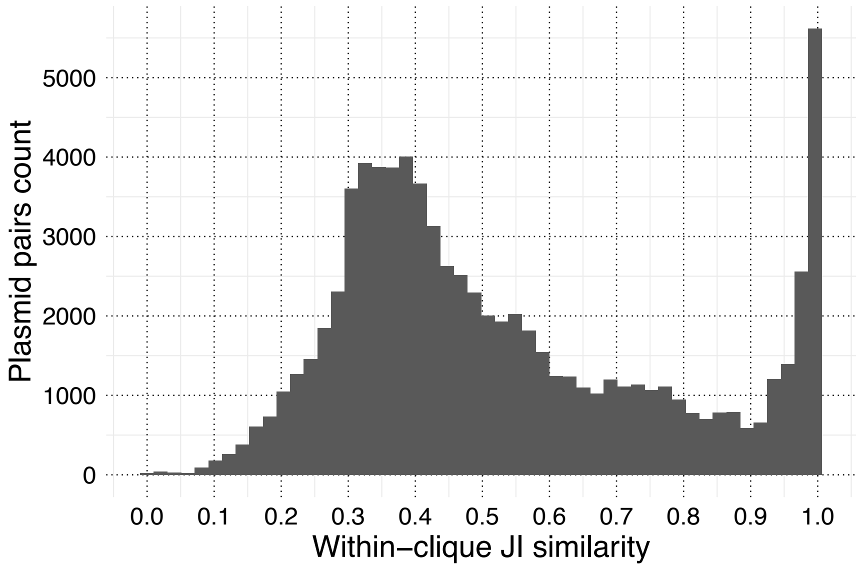


**Supplementary Figure 7. The distribution of Jaccard Index (JI) similarities between plasmid pairs within cliques.**

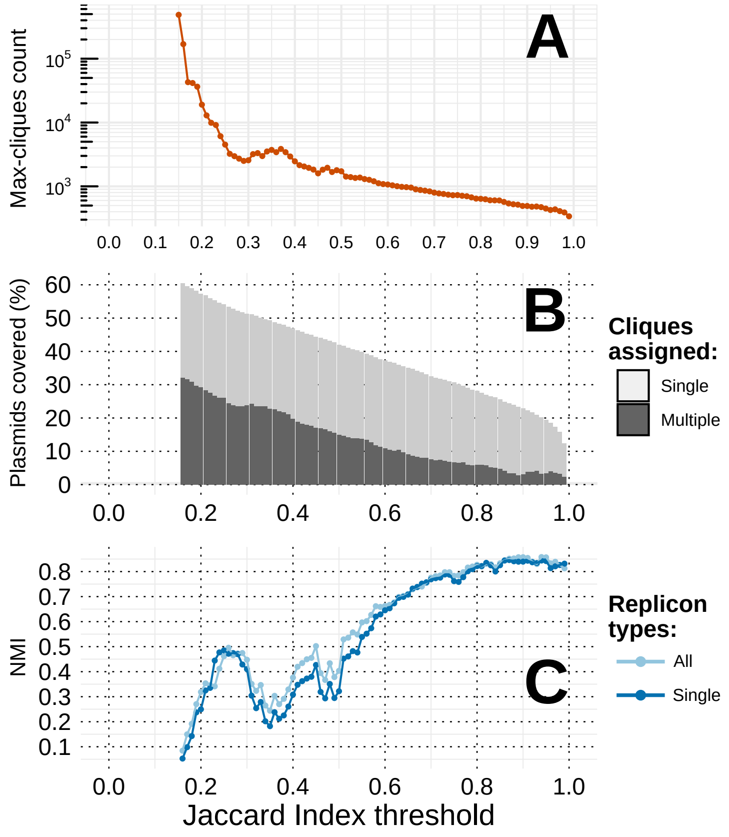


**Supplementary Figure 8. Assesment of the Max-clique algorithm performance over a range of Jaccard Index (JI) thresholds.** JI thresholds were used to transform the weighted plasmid network (Supplementary Figure 3) to a binary one after which a Max-clique algorithm was used to identify all maximal cliques of size three or more. For each JI value, the results presented here show: total number of maximal cliques detected (A), percentage of plasmids covered by the cliques (B), and the congruence with replicon typing measured by NMI score (C). The analysis below JI of 0.15 was aborted due to high memory and computational requirements due to the large number of detected cliques.

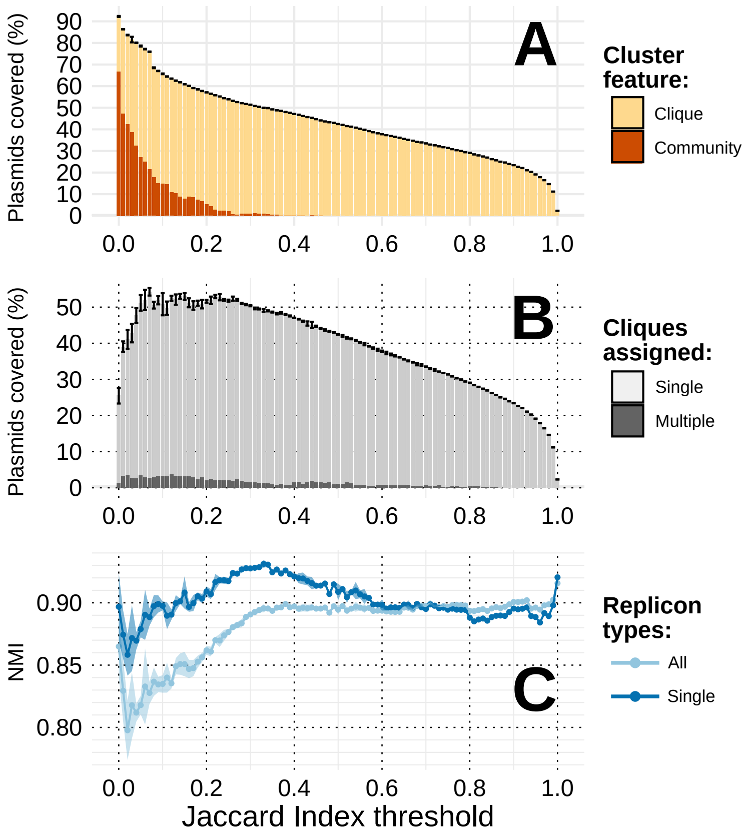


**Supplementary Figure 9. OSLOM performance over a range of Jaccard Index (JI) thresholds after the removal of 29,913 accessory CDSs from the plasmid sequences.** Similarly to main text Figure 2, edges with values below a particular JI threshold were removed from the original plasmid network (Supplementary Figure 3) prior to the OSLOM analysis. The OSLOM performance was assessed based on the following criteria: (A) clique to community ratio; (B) percentage of plasmids covered by the cliques; (C) the congruence with replicon typing measured by NMI score. Error bars (A and B) and light-coloured shading (C) provide +/- 2 standard deviations (SD) of uncertainty. Standard deviation around every value on the y-axis across all JI thresholds assessed (points and bars) was calculated based on results of n=5 iterations of OSLOM software (see Methods). Maximal NMI score detected was 0.9157 for plasmids assigned to a single or multiple replicon types at JI=1.0, and 0.9312 for plasmids belonging to only one replicon type at JI=0.33.

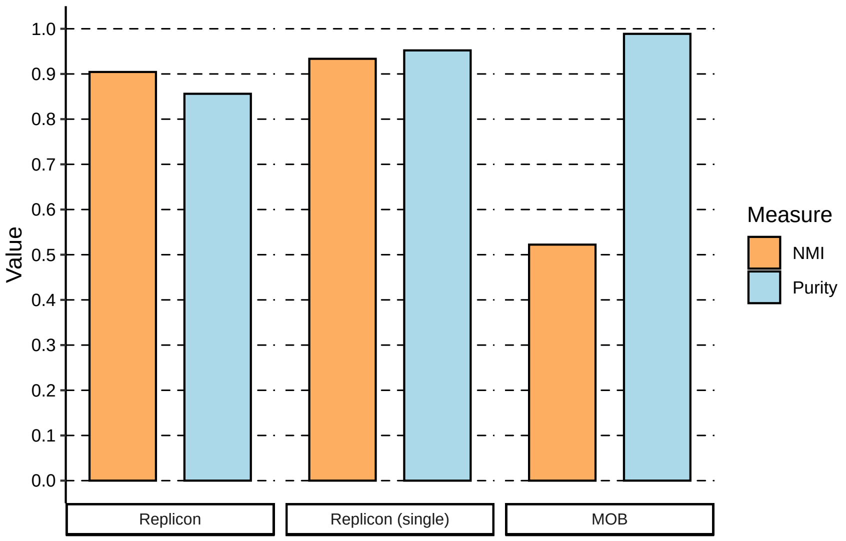


**Supplementary Figure 10. Concordance of plasmid clique assignment with replicon and MOB typing schemes.** Normalized Mutual Information (NMI) and purity (see Methods) were calculated for cliques containing plasmids with identified replicon type, plasmids carrying a single identified replicon type, and plasmids assigned to MOB types.

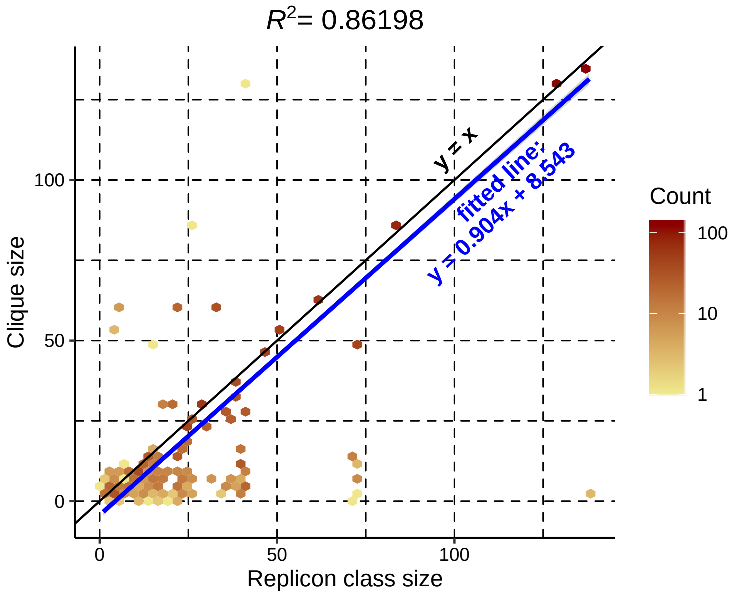


**Supplementary Figure 11. Plasmid clique size as a function of replicon class size.** For each single-type plasmid in the dataset, the sizes of its corresponding clique and replicon class size were determined, y and x axis respectively. The colour intensity of each hexagon reflects the density of the data points (count scale). If all plasmids from a particular replicon type are encompassed by a single clique, the points corresponding to those plasmids would fall on the line $y=x$. The coefficient of determination ($R^2$) for the function $y=x$ was estimated to be 0.86198. In addition, the slope of the function ($y=0.904x + 8.543$) derived from the data points by linear regression is less than that of a $y=x$ which reflects the trend of plasmids from the same replicon class being sorted into multiple smaller cliques.

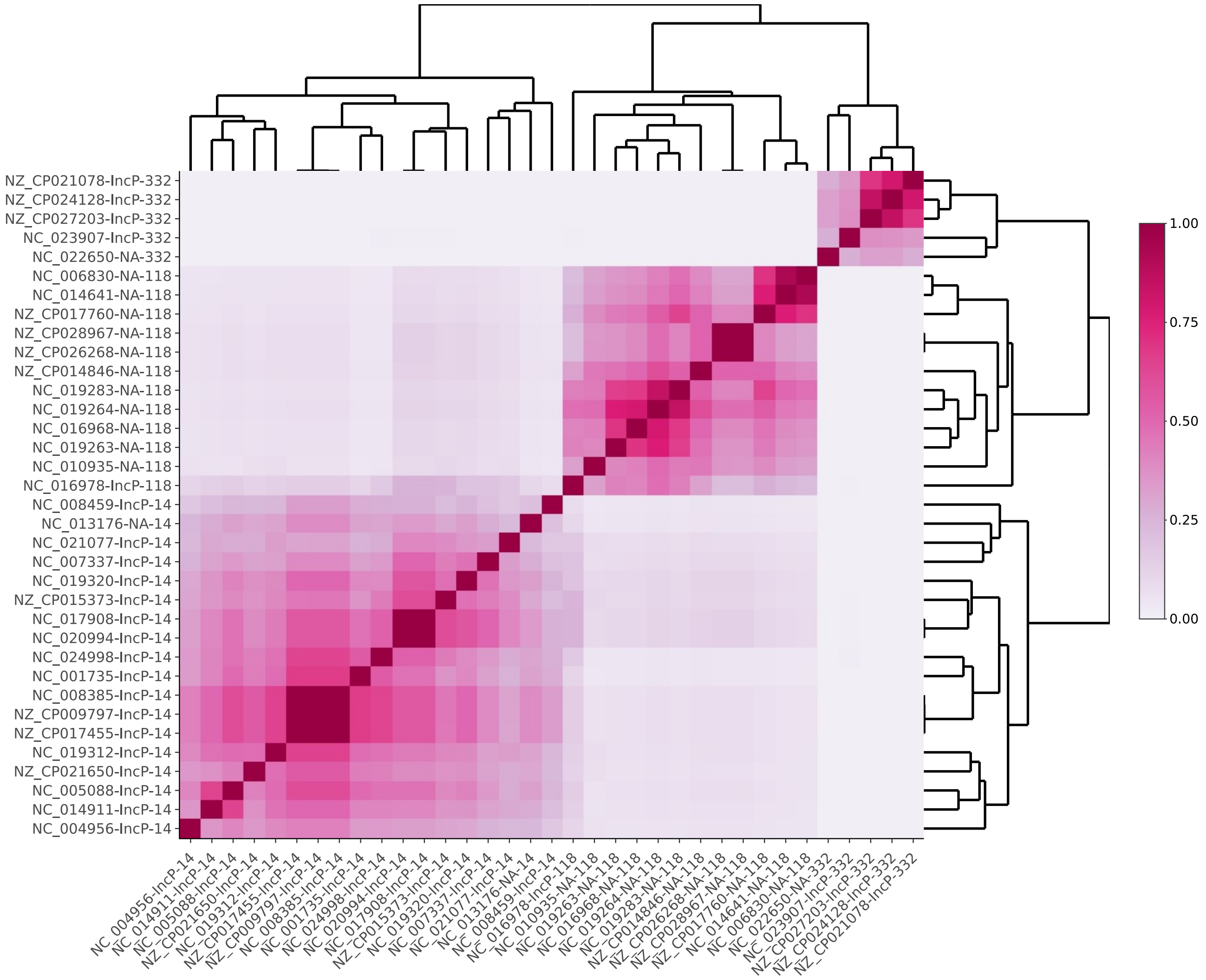


**Supplementary Figure 12. Heatmap of pairwise Jaccard Index (JI) distances between plasmids from cliques containing IncP replicon type.** Plasmids were ordered using hierarchical clustering as provided by the two dendrograms. The legend on the right matches the colour gradient of the heatmap with the corresponding JI value. The accession number of each plasmid sequence can be found on two symmetrical axes and it is followed by the replicon type and the clique number. NA denotes plasmids with unknown replicon type.

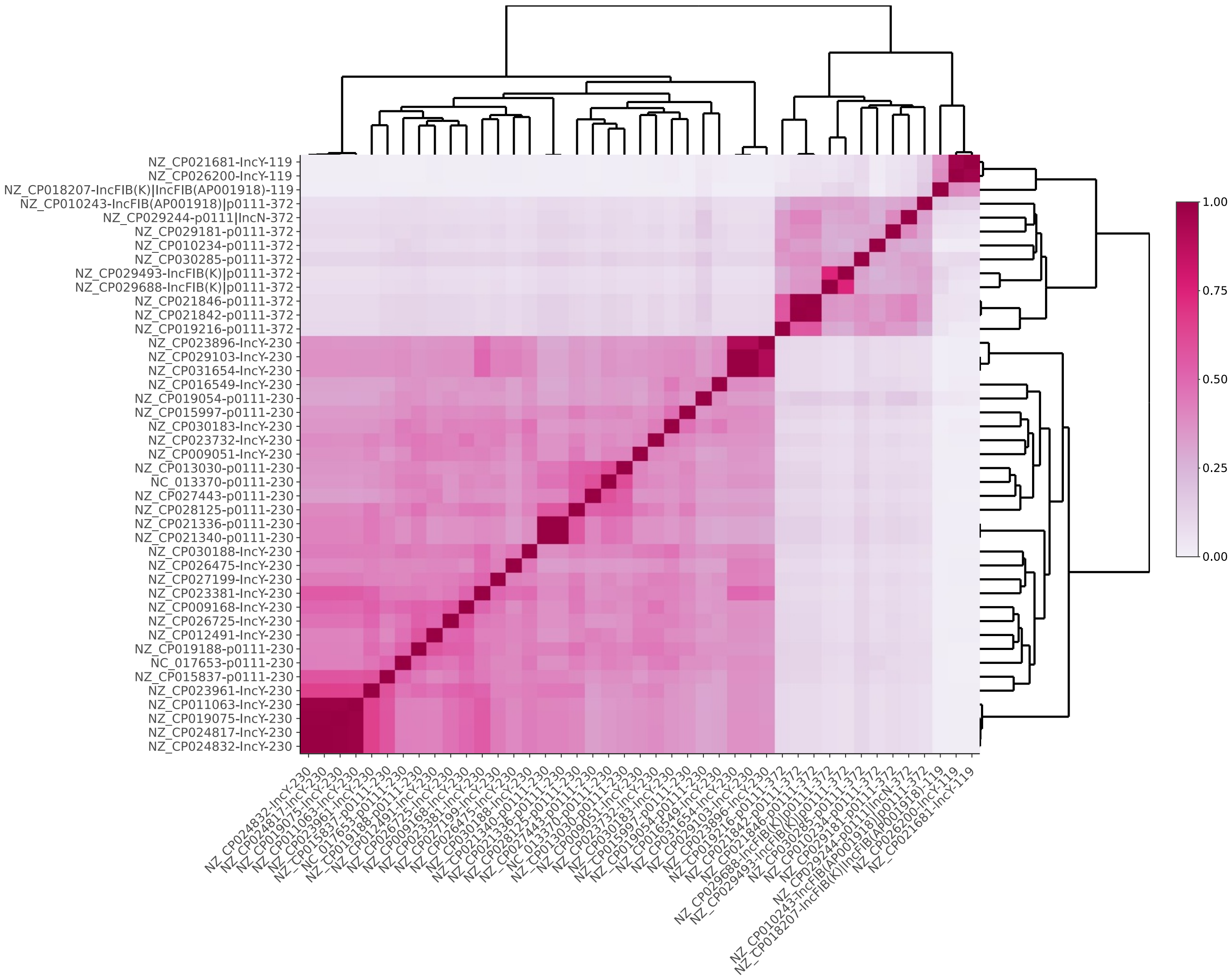


**Supplementary Figure 13. Heatmap of pairwise Jaccard Index (JI) distances between plasmids from cliques containing IncY and p0111 replicon types.** Plasmids were ordered using hierarchical clustering and their relatedness is provided by the two dendrograms. The legend on the right matches the colour gradient of the heatmap with the corresponding JI value. The accession number of each plasmid sequence can be found on two symmetrical axes and it is followed by the replicon type and the clique number.

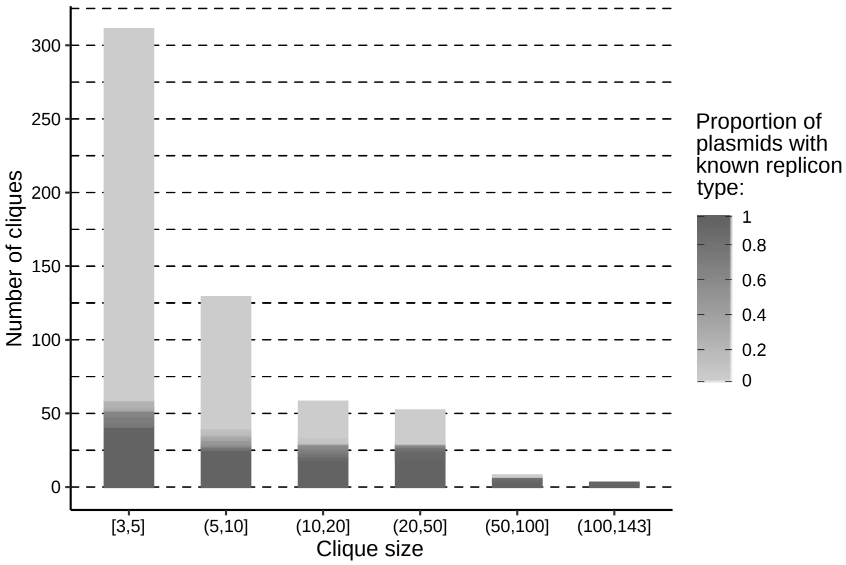


**Supplementary Figure 14. The distribution of the clique sizes.** The cliques were sorted into six bins based on the number of plasmids they carry (*x*-axis). The shading on each bin corresponds to the number of cliques within a bin that have a certain proportion of plasmids with known replicon type (as indicated by the figure legend at right).

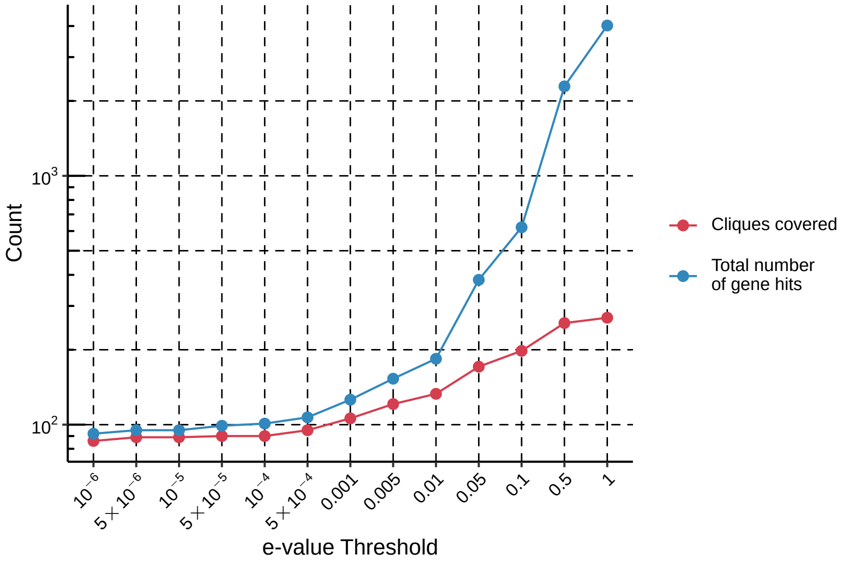


**Supplementary Figure 15. Finding the optimal e-value threshold for discovery of candidate replicon genes within untyped plasmid cliques.** The core genes of the untyped cliques were screened against the PlasmidFinder replicon database using TBLASTN. The number of cliques covered by the blast search and the total number of core gene hits were recorded. The majority of plasmids within the dataset are assigned to a single replicon type though with some plasmids assigned to a maximum of three to four different types. Upon reaching the e-value of 0.01, the number of gene hits starts rapidly diverging from the number of cliques covered disclosing the false positives. Therefore, a conservative e-value of 0.001 was chosen as the final threshold.

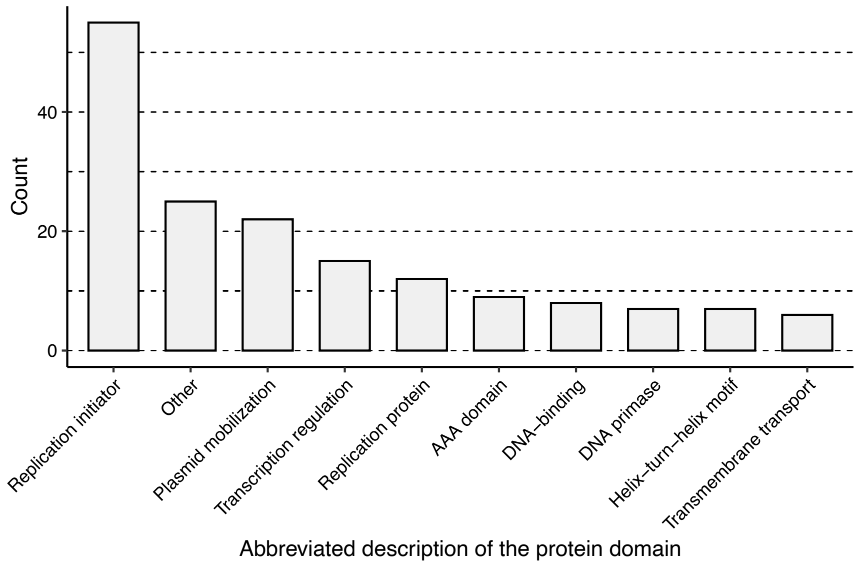


**Supplementary Figure 16. Protein domain families found associated with the candidate replicon genes.** Protein sequences of candidate replicon genes were screened using HMMER tool against the Pfam database. The domain families discovered were binned according to their abbreviated description.

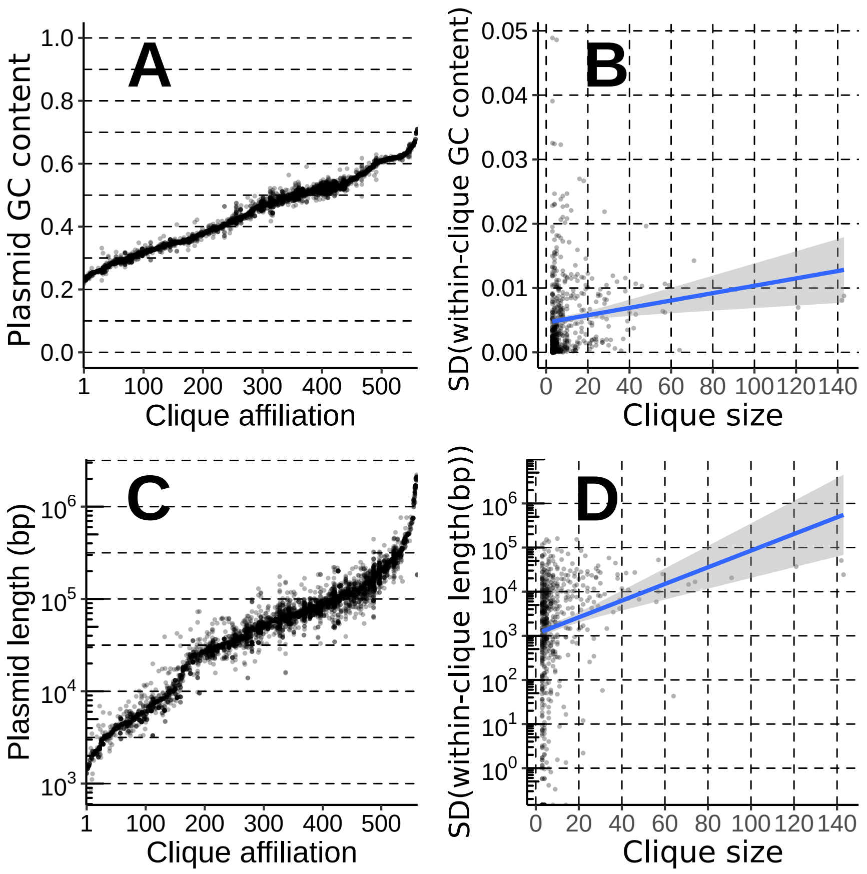


**Supplementary Figure 17. Variability of plasmids within cliques in GC content and length.** The GC content (A) and the length (C) of a particular plasmid (*y*-axis) relative to its clique affiliation (*x*-axis columns). The clique affiliation of plasmids has been ordered based on the average clique GC content and the average clique length in panels A and C respectively. Hence, the numbers on both x-axes in panels A and C are arbitrary and do not correspond to the same clique. To further explore within-clique variability, the standard deviation (SD) of the within-clique GC content (B) and length (D) were considered with respect to the clique size. A weak linear correlation with clique size was found for both SD measurements ($R^2$ = 0.0155 and $R^2$ = 0.029 for B and D respectively).

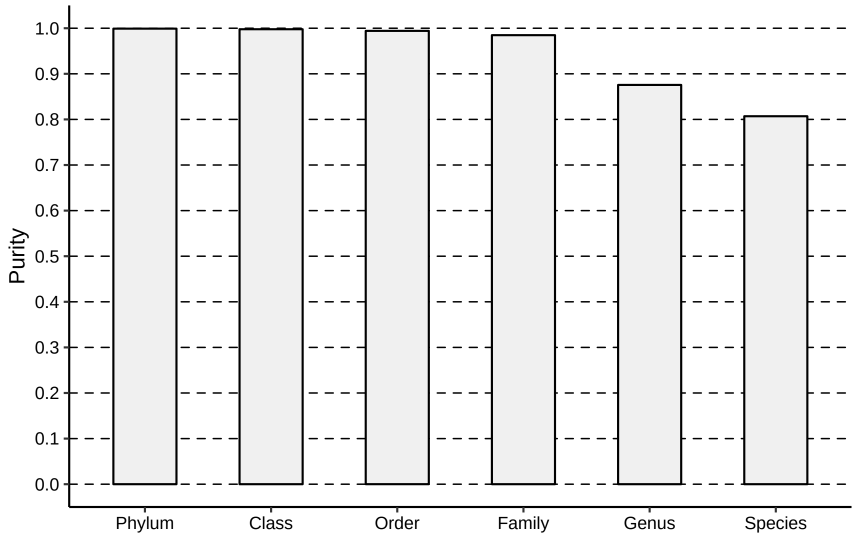


**Supplementary Figure 18. Purity of cliques relative to the taxonomic level of the plasmid bacterial host.**

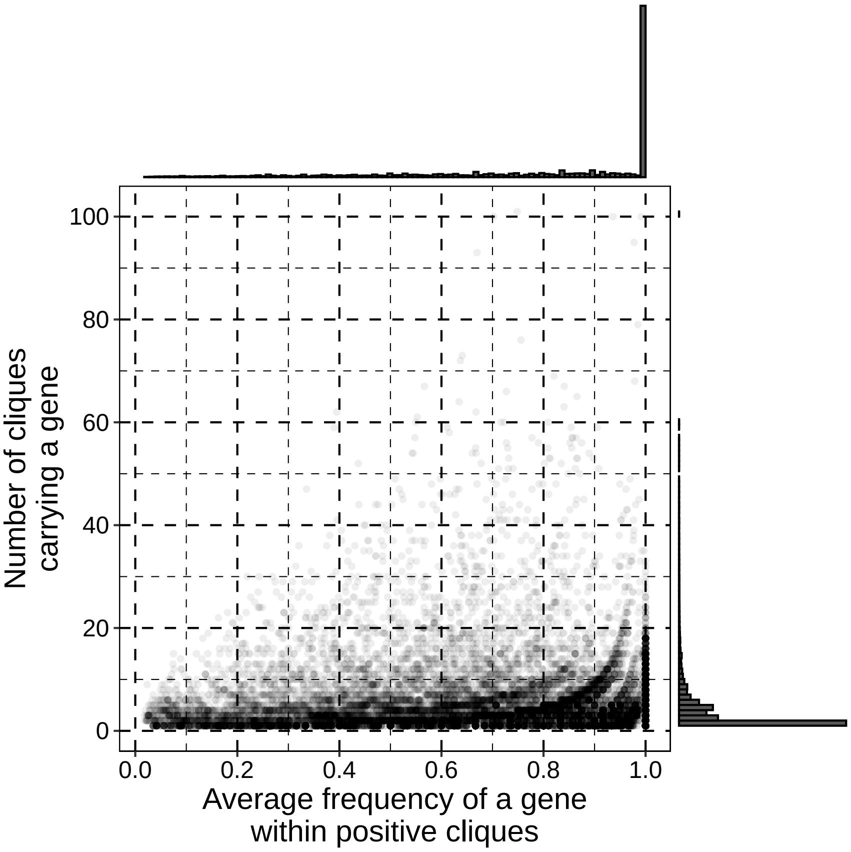


**Supplementary Figure 19. Assessing the frequency of genes within cliques.** The average within-clique frequency of all genes with five or more occurrences in the dataset was calculated and plotted against the number of cliques in which a particular gene occurs. Within-plasmid gene duplications were disregarded and counted as a single occurrence. The histograms on the top and the right-hand side provide the distribution of values for the two axes.

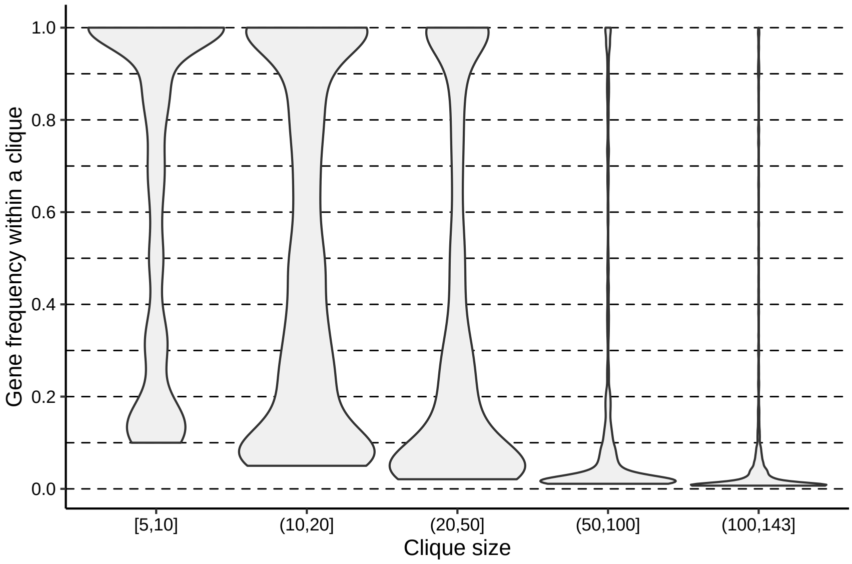


**Supplementary Figure 20. The distribution of within-clique gene frequencies relative to the clique size.** The assessed genes had five or more occurrences in the dataset, thus only the cliques carrying five or more plasmids were considered.

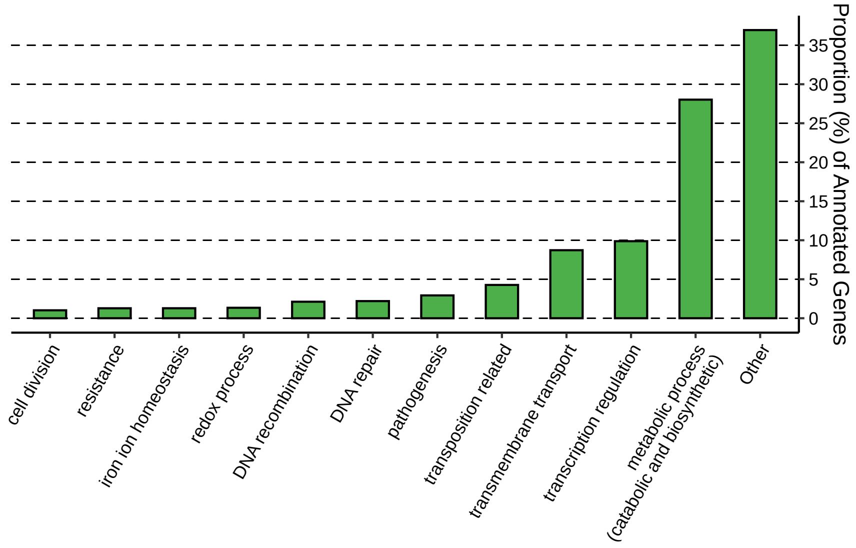


**Supplementary Figure 21. Distribution of the biological functions associated with the core genes in plasmid cliques.** The respective frequency (%) of each gene was considered.

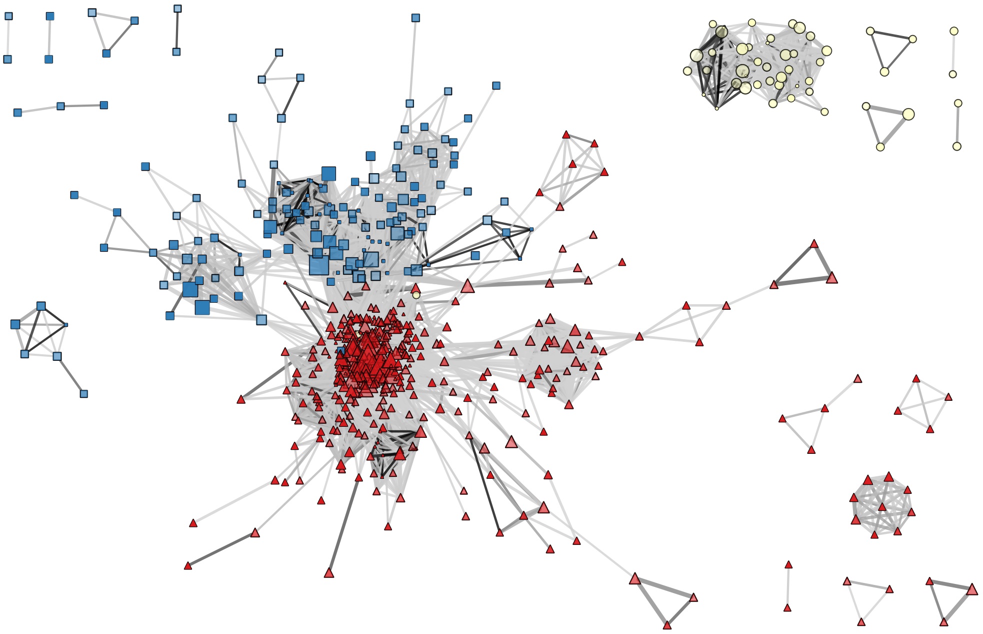

**Supplementary Figure 22. The unfiltered network of plasmid cliques.** Colour and shape of the cliques (vertices) indicate the phylum of the predominant bacterial host: Proteobacteria – red triangle; Firmicutes – blue squares; Other phyla – yellow circles. As noted in the legend of main text Figure 4, the transparency of the vertex indicates the average internal JI of the clique, the colour of the edges indicates the average JI between plasmids of two cliques, and the width of the edge is proportional to the number of connections.
